# Dissecting HIV Virulence: Heritability of Setpoint Viral Load, CD4+ T Cell Decline and Per-Parasite Pathogenicity

**DOI:** 10.1101/140012

**Authors:** Frederic Bertels, Alex Marzel, Gabriel Leventhal, Venelin Mitov, Jacques Fellay, Huldrych F Günthard, Jürg Böni, Sabine Yerly, Thomas Klimkait, Vincent Aubert, Manuel Battegay, Andri Rauch, Matthias Cavassini, Alexandra Calmy, Enos Bernasconi, Patrick Schmid, Alexandra U Scherrer, Viktor Müller, Sebastian Bonhoeffer, Roger Kouyos, Roland R Regoes, the Swiss HIV Cohort Study

## Abstract

Pathogen strains may differ in virulence because they attain different loads in their hosts, or because they induce different disease-causing mechanisms independent of their load. In evolutionary ecology, the latter is referred to as “per-parasite pathogenicity”. Using viral load and CD4+ T cell measures from 2014 HIV-1 subtype B infected individuals enrolled in the Swiss HIV Cohort Study, we investigated if virulence — measured as the rate of decline of CD4+ T cells — and per-parasite pathogenicity are heritable from donor to recipient. We estimated heritability by donor-recipient regressions applied to 196 previously identified transmission pairs, and by phylogenetic mixed models applied to a phylogenetic tree inferred from HIV *pol* sequences. Regressing the CD4+ T cell declines and per-parasite pathogenicities of the transmission pairs did not yield heritability estimates significantly different from zero. With the phylogenetic mixed model, however, our best estimate for the heritability of the CD4+ T cell decline is 17% (5%–30%), and that of the per-parasite pathogenicity is 17% (4%–29%). Further, we confirm that the set-point viral load is heritable, and estimate a heritability of 29% (12%–46%). Interestingly, the pattern of evolution of all these traits differs significantly from neutrality, and is most consistent with stabilizing selection for the set-point viral load, and with directional selection for the CD4+ T cell decline and the per-parasite pathogenicity. Our analysis shows that the viral genetype affects virulence mainly by modulating the per-parasite pathogenicity, while the indirect effect via the set-point viral load is minor.

## 1 Introduction

The virulence of an infection is determined by both, the host and the pathogen. One of the most common modulators of virulence is the pathogen load. Higher load often leads to more morbidity, or to faster disease progression or death. Similar to virulence, the load that a pathogen strain attains is also determined by both, the host and the pathogen. In evolutionary ecology, hosts that limit virulence by reducing pathogen load are called “resistant”, and pathogen strains that attain a high load in their hosts are often termed “virulent”.

But virulence is not completely determined by the pathogen’s load alone. There are pathogen-load-independent components, which are again influenced by the host and the pathogen. A host that suffers less than average from being infected by a pathogen and carrying a specific load is called “tolerant” (Ayres and Schneider, 2012; Boots, 2008; Boots *et al*., 2009; Caldwell *et al*., 1958; Little *et al*., 2010; Medzhitov *et al*., 2012; Råberg, 2014; Råberg *et al*., 2007, 2009; Read *et al*., 2008; Schafer, 1971; Schneider and Ayres, 2008). A pathogen strain that causes less than average virulence attaining a specific load is said to have a low “per-parasite pathogenicity” (Råberg, 2014; Råberg and Stjernman, 2012). Fig 1A displays these virulence components diagrammatically.

**Figure 1:**
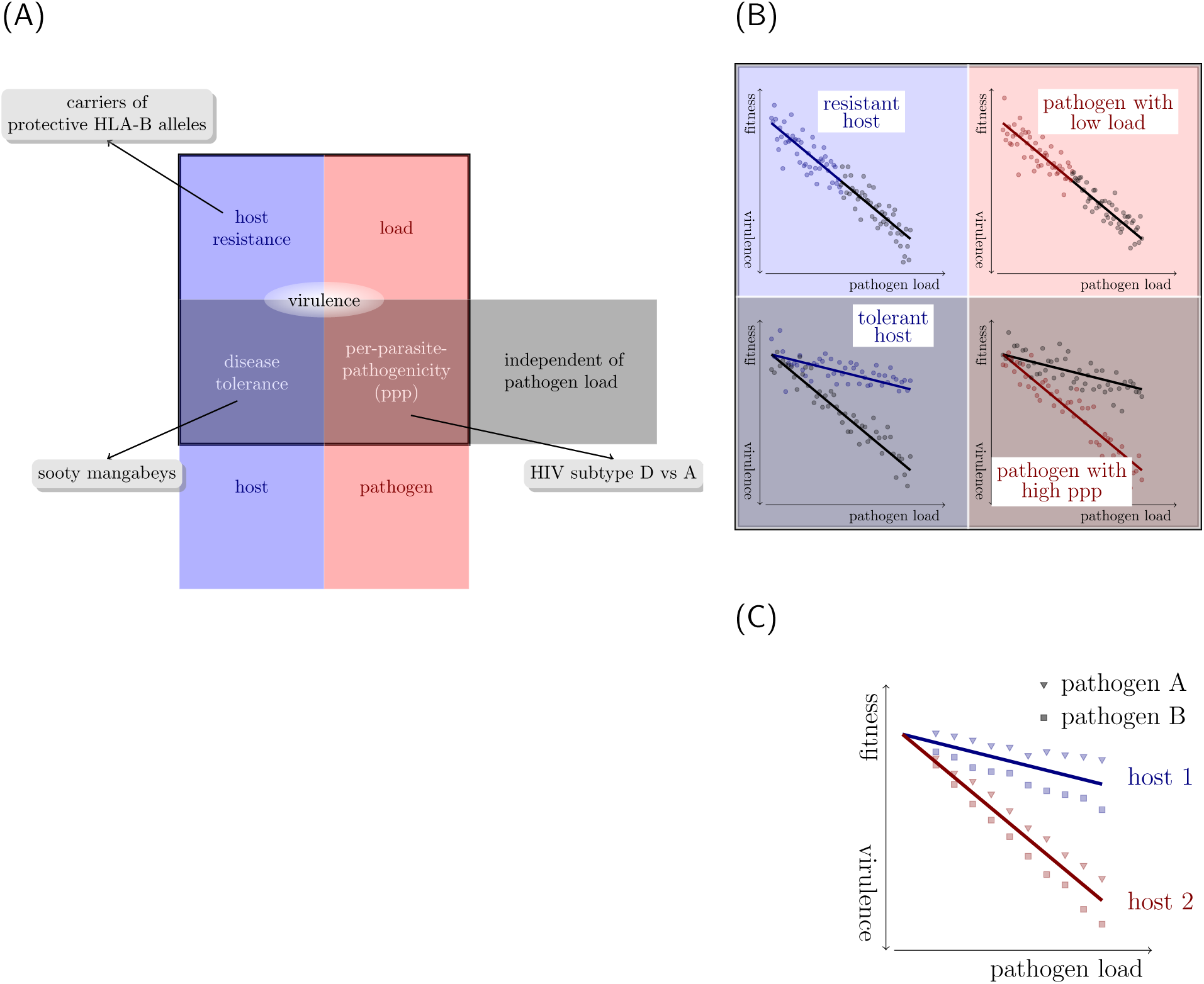
Dissecting virulence. (A) Systematics of virulence components. Each component can be a trait of either the pathogen or the host, and depend or be indepependent of the load of the pathogen. (B) Formally, virulence can be dissected using fitness-versus-pathogen-load plots. (In these plots, host fitness is inversely correlated with virulence.) Adapted from Figure 1 in Raberg (Råberg, 2014). (C) In multi-host multi-pathogen systems, virulence components can be disentangled by first defining host-type-specific tolerance curves. Pathogens differing in their per-parasite pathogenicity will then fall on different sides of these tolerance curves. In the example shown, pathogen B has a higher per-parasite pathogenicity than pathogen A.

How can pathogen-load-independent components of virulence be determined? To identify these components, “excess virulence” needs to be measured, i. e. by how much virulence differs from what is expected for a specific pathogen load. Statistically speaking, “excess virulence” is the residual virulence after adjusting for differences in pathogen load. This adjustment can be visualized on fitness-versus-pathogen-load plots (Fig 1B). On such a plot, host types with differing levels of disease tolerance are characterized by different tolerance curves that depict the relationship between virulence and pathogen load (see Fig 1B bottom left). The steepness of this curve is inversely related to disease tolerance.

Once tolerance curves for different host types are determined, the per-parasite pathogenicity will manifest itself as a yet unexplained deviation from the tolerance curve. In other words, varying degrees of per-parasite pathogenicity will lead to residual excess virulence that is not explained by host factors. Fig 1C shows how two pathogen strains differing in their per-parasite pathogenicity will scatter around the tolerance curves of two host types.

HIV infection provides an ideal example to illustrate this decomposition of virulence. In this infection, CD4+ T cells — the target cells of the virus — continuously decline from a level of approximately 1000 cells per microliter blood. A CD4+ T cell count below 200 cells per microliter blood is one of the defining characteristics of AIDS. The decline rate of the CD4+ T cells is a well-established surrogate for the rate of disease progression (Phillips *et al*., 1991), i.e. virulence. It has the advantage that it can be determined from clinical samples spanning less than one year of monitoring an HIV infected individual, while the direct observation of disease progression requires many years. The existence of such a well-established, quantitative measure of virulence makes HIV infection unique.

The decomposition of virulence relies on its relation with pathogen load. During HIV infection, the viral load rises and peaks a few weeks after infection, and subsequently settles at a fairly stable level that is maintained for many years — the so-called set-point viral load. This set-point viral load represents a good measure of pathogen load required for the decomposition of virulence, and it is associated with the rate of progression toward disease and death (Mellors *et al*., 1996). However, the correlation between the set-point viral load and the decline of the CD4+ T cells — a good proxy of the rate of disease progression — is not very strong: *R*^2^ values were found to be between 0.05–0.08 in American cohorts (Rodriguez *et al*., 2006), and 0.05 for the population studied here (Regoes *et al*., 2014) (although the relationship between set-point viral load and survival time has been reported to be higher (Arnaout *et al*., 1999).) While this weak correlation may be in part the result of measurement error it suggests that there are factors influencing virulence other than the set-point viral load.

In the context of HIV infection, examples for variation in all of the four virulence components exist. First, the set-point viral load differs by three orders of magnitude between HIV infected individuals, ranging from 10^3^ to 10^6^ RNA copies per ml plasma (Deeks *et al*., 1997; Mellors *et al*., 1996; Raboud *et al*., 1996). This variation is associated with the rate of disease progression (Mellors *et al*., 1996), i.e. virulence. Recent heritability studies (as reviewed in Fraser *et al*. (2014)) have shown that a fraction of the variation in the set-point viral load can be attributed to the viral genotype infecting the host.

Second, human genes conferring “host resistance” in the sense of evolutionary ecology (see above) have been identified: Individuals who carry protective HLA-B alleles have lower set-point viral loads, and progress to disease at a slower rate than people without these alleles (Fellay *et al*., 2007; Goulder and Watkins, 2008). Across primates, species-specific restriction factors can limit the load of Human and Simian Immunodeficiency Viruses (Kirchhoff, 2009; Zheng *et al*., 2012).

Third, in the context of immunodeficiency viruses, examples for variation in pathogenload-independent virulence components exist. Simian Immunodeficiency Virus (SIV) infection in natural hosts, such as the sooty mangabeys, is avirulent despite the high set-point viral loads SIV attains in these hosts (Chahroudi *et al*., 2012; Chakrabarti, 2004). In contrast, SIV infection in non-natural hosts, such as the rhesus macaque, leads to an AIDS-like disease (Chahroudi *et al*., 2012; Chakrabarti, 2004). Thus, natural hosts tolerate the infection without becoming sick. There is also variation in tolerance to HIV infection in humans associated with age and human leukocyte antigens (HLA) (Regoes *et al*., 2014).

Lastly, there is also variation in per-parasite pathogenicity across viral strains. For example, there is evidence that HIV-1 subtype D leads to faster disease progression than subtype A even though both subtypes attain similar set-point viral loads (Baeten *et al*., 2007). In other words, subtype A and D differ in their per-parasite pathogenicity.

Previously, we investigated if humans display variation in disease tolerance against HIV (Regoes *et al*., 2014). We found that younger individuals and HLA-B heterozygotes are more tolerant, and could link the variation in tolerance to HLA-B genotype. In this previous study, we did, however, not investigate the potential impact of the virus genotype on virulence.

How can one investigate if the virus genotype influences a host-pathogen trait? One way is to study associations between genetic polymorphisms of the virus and the trait, as is done, for example, in genome-wide association studies (Bartha *et al*., 2013). An alternative approach is to estimate the heritability of the trait. If a trait is heritable, i. e. similar between similar viral genotypes, it must, at least in part, be determined by viral genes.

The influence of the virus genotype on the set-point viral load has been the focus of many research groups, including ours, over the past years. Most studies determined the heritability of the set-point viral load (Alizon *et al*., 2010; Fraser *et al*., 2014; Hodcroft *et al*., 2014; Hollingsworth *et al*., 2010; Müller *et al*., 2011), while others investigated associations between genetic polymorphisms of the virus and the trait (Bartha *et al*., 2013). There is a consensus that set-point viral load is heritable, although there is some controversy on the numerical value of the heritability.

In this study, we investigate the influence of the HIV genotype on overall virulence, as measured by the CD4+ T cell decline, and its pathogen-load-dependent and-independent components, the set-point viral load and per-parasite pathogenicity, respectively. We determine the influence of the viral genotype by estimating the “heritability” of these traits, i. e. the correlation of each of these traits in donor and recipient. To this end, we use data from the Swiss HIV Cohort Study. For our analysis, we selected cohort participants, for whom we could determine the set-point viral load and the decline of CD4+ T lymphocytes — an established proxy for virulence in HIV infection. As a surrogate for the per-parasite pathogenicity we use the residuals from previously determined tolerance curves (Regoes *et al*., 2014) as described in Methods and Materials.

Our analysis confirms that set-point viral load is heritable. We further provide evidence that the decline of CD4+ T lymphocytes, i. e. virulence, is heritable. Lastly, we find evidence for the heritability of the per-parasite pathogenicity, the pathogenload-independent component of virulence. Our results are therefore consistent with the notion that the virus genotype affects virulence in HIV infection both via the viral load, and via viral-load-independent mechanisms.

## 2 Methods and Materials

### 2.1 Study population

We analysed a subset of the individuals from the Swiss HIV Cohort Study (www.shcs.ch) (Schoeni-Affolter *et al*., 2010). This study has enrolled more than 19’000 HIV-infected individuals to date, which constitutes more than 72% of all patients receiving antiretroviral therapy in Switzerland, and is therefore highly representative. The viral load and CD4+ T cell count of each enrolled individual are determined approximately every three months. In some of these individuals, the *pol* gene of the virus was sequenced. The *pol* gene encodes viral enzymes important for viral replication within its target cell, most notably the reverse transcriptase, the integrase, and the protease, and contains many of the clinically relevant resistance mutations.

The study population of the present study consists of a subset of the study population analyzed in a previous study (Regoes *et al*., 2014). In this previous study, we had included 3036 HIV-1 infected individuals, for whom viral load measurements and CD4+ T cell counts were available to reliably estimate the set-point viral load and CD4+ T cell decline. We restricted our analysis to data obtained before antiretroviral treatment. Furthermore, we excluded the primary and late phases of the infection by discarding measurements during the first 90 days after the estimated date of infection and measurements obtained when the CD4+ T cell count was below 100 per microliter blood. Lastly, individuals were included if they had at least two viral load measurements and three CD4+ T cell measurements that were at least 180 days apart.

For the present study, we selected 2014 individuals of the 3036 individuals enrolled previously. Individuals were included if the *pol* gene of their virus had been sequenced. The genetic information of the virus was necessary for the present study to infer the evolutionary history and investigate patterns of heritability.

*Pol* sequence information was obtained from the SHCS genotypic drug resistance database. Sequences are stored in a central database (SmartGene; Integrated Database Network System version 3.6.13). All laboratories perform population-based sequencing (von Wyl *et al*., 2007; Yang *et al*., 2015). The drug resistance database includes, in addition to the routinely collected samples, over 11’000 samples from the biobank analyzed by systematic retrospective sequencing (Schoeni-Affolter *et al*., 2010; Yang *et al*., 2015). The individuals in our study population belong to the following risk groups: men having sex with men – 972 (48%), heterosexuals – 435 (21.5%), intravenous drug users – 365 (18%), and others – 252 (12.5%).

The SHCS, enrolling HIV-infected adults aged over 16 years old, has been approved by ethics committees of all participating institutions. The data collection was anonymous and written informed consent was obtained from all participants (Schoeni-Affolter *et al*., 2010).

### 2.2 Set-point viral loads and CD4+ T cell declines

For each individual enrolled in our study, the set-point viral load had been determined in a previous study (Regoes *et al*., 2014) as the mean of the logarithm to the base 10 of the eligible viral load measurements in each individual. Non-detectable viral loads had been set to half the detection limit. The rate at which the CD4+ T cells change per day had previously been estimated as the slope in a linear regression of CD4+ T cell counts in an individual against the date, at which they were determined. The rate of change in CD4+ T cells is inversely related to virulence.

Set-point viral loads and CD4+ T cell declines were adjusted for potential covariates by regressing them against sex, age at infection, risk group and ethnicity. Once significant covariates were identified, adjusted traits were defined as the residuals of a regression with these covariates. Subsequent analyses were then conducted with the residuals.

The inclusion criteria, calculation of set-point viral load and CD4+ T cell decline, as well as the model fitting and comparisons had been implemented and performed in the R language of statistical computing (R Core Team, 2013).

### 2.3 Per-parasite pathogenicity

Per-parasite pathogenicity is defined as the pathogenic potential of a pathogen strain adjusted for its load and host factors that are associated with pathogen-load independent virulence components, i.e. tolerance.

To derive a proxy for the per-parasite pathogenicity of a strain, we first determined the relationship between pathogen load and pathogenicity — called the tolerance curve — for a given host type. In our previous study (Regoes *et al*., 2014), we found that the age, at which the host was infected, was associated very strongly with the slope of the tolerance curve. Therefore, we determined the tolerance curve specific for the age of the host that harbors the pathogen strain. We did not account for host factors other than age, such as, for example, HLA-B genotype or homozygosity, because this information of host genotype is lacking for the majority of our study population.

In a next step, we predicted the CD4+ T cell decline we should observe given the set-point viral load that this strain attains in the host. Lastly, we calculated by how much the observed CD4+ T cell decline deviates from this prediction (see Fig 2).

**Figure 2:**
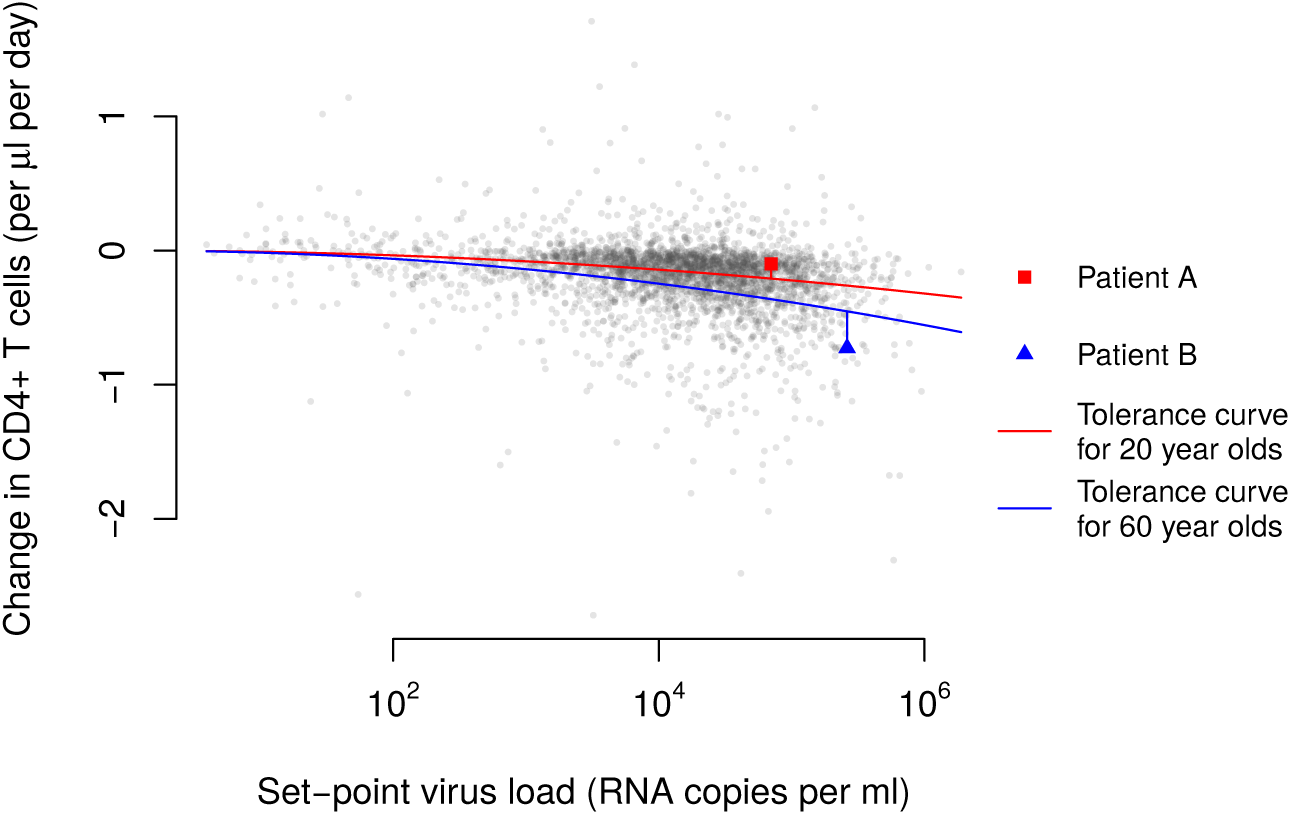
As a surrogate for the per-parasite pathogenicity we use the residuals from age-adjusted tolerance curves. In the graph, we plotted each individual’s CD4+ T cell decline versus his/her set-point viral load. The red and blue curves show the average relationships between these two measures in the groups that were 20 and 60 years old at the time of their infection, respectively. These age-adjusted tolerance curves were determined previously (Regoes *et al*., 2014). The red square and blue triangle highlight two individuals with an age at infection of 20 and 60 years, respectively.

In our conceptualization, tolerance is measured by the parameter characterizing the relationship between the CD4+ T cell decline and the set-point viral load. Different levels of per-parasite pathogenicity, on the other hand, manifest themselves as a deviation of an individual’s CD4+ T cell decline and set-point viral load from the tolerance curve. Formally, this procedure amounts to regressing CD4+ T cell decline against the setpoint viral load, adjusting for the age at infection, and calculating the residual of an individual’s trait from the regression line. We denote this proxy as ppp. Lower and more negative values of this quantity are associated with faster disease progression.

### 2.4 Transmission pairs and phylogenetic tree

The transmission pairs were identified as monophyletic clusters on a previous HIV transmission tree (Kouyos *et al*., 2014). Of these previously established transmission pairs, 196 were present in our study population. The direction of transmission cannot be inferred in these pairs.

The reconstruction of the phylogenetic tree relies on *pol* gene sequencing of the virus carried by the study subjects. In particular, we had sequences of pol extending over the HXB2 positions 2253–3870, comprising the protease and the reverse transcriptase. All sequences were initially aligned to an HXB2 reference genome (http://www.ncbi.nlm.nih.gov/nuccore/K03455.1) using MUSCLE (Edgar, 2004). We selected the earliest sequence if more than one sequence was available for a person.

To reconstruct the evolutionary history, we first removed insertions relatively to HXB2. To exclude signatures of parallel evolution due to drug pressure that can distort the inferred evolutionary history, we further removed drug resistance mutations according to the databases of Stanford (http://hivdb.stanford.edu/) and the International Antiviral Society (https://www.iasusa.org/). We used Gblocks to refine the alignment. The final number of positions was 1106.

We constructed the phylogenetic tree using FastTree (Version 2.1.8 SSE3, OpenMP) (Price *et al*., 2010). We used a maximum-likelihood-based inference using a Generalized Time-Reversible evolutionary model and a CAT model (Stamatakis, 2006a) with 20 discrete evolutionary rate categories. We use the most rigorous and time-consuming FastTree parameters (FastTreeMP-pseudo-spr 4-mlacc 2-slownni-gtr-nt). We rooted the tree with 10 Subtype C sequences as an outgroup, using the R package APE. The branch lengths in our tree correspond to genetic distances between the sequences, and not to time.

We also compared the results obtained from this tree with those of two trees reconstructed with RaxML (Stamatakis, 2006b). These trees were reconstructed assuming a Generalized Time-Reversible evolutionary model. One assumed CAT model with 25 discrete evolutionary rate categories, the other Γ-distributed evolutionary rates. Table S1 shows that there is genrally good agreement between the results.

### 2.5 Heritability estimation

To estimate the heritability of the three traits — set-point viral load, CD4+ T cell decline and per-parasite pathogenicity — we used two approaches.

First, we applied donor-recipient regressions that are formally equivalent to parent-offspring regressions (Fraser *et al*., 2014) to the 196 previously identified transmission pairs. Although we do not have any information on the direction of transmission in these pairs, we expect that a regression between the trait in question will yield a good estimate of the heritability (Bachmann *et al*., 2017).

Second, we employed phylogenetic mixed models (Housworth *et al*., 2004) that are widely used to estimate the heritability from a phylogenetic tree. These methods have the advantage of being able to incorporate larger study populations and transmission relationships ranging from close pairing to distant epidemiological linkage. We used the recent implementation by Leventhal and Bonhoeffer (Leventhal and Bonhoeffer, 2016) in the R language of statistical computing (R Core Team, 2013). The models that underlie this method assume trait evolution according to Brownian motion, i. e. that traits drift neutrally. Brownian motion is characterized by the diffusion constant which is related to the heritability. Because we use the method on a tree, the branch lengths of which correspond to genetic distances, we make the implicit assumption that heritability increases linearly with genetic distance. Previous studies used Pagel’s λ to estimate heritabilities (Alizon *et al*., 2010; Shirreff *et al*., 2013). We refrained from using Pagels λ in our main analysis because it is not as appropriate as phylogenetic mixed model for non-ultrametric trees (as our phylogenetic tree) (Leventhal and Bonhoeffer, 2016). However, as a point of comparison with estimates of Pagels λ in previous studies, we report our estimates of this quantity in the supplementary material.

We also applied phylogenetic mixed models based on the Ornstein-Uhlenbeck process that describe stabilizing trait selection around an optimal trait value rather than neutral drift. In addition to the diffusion constant, this model has a parameter for the trait value around which selection stabilizes the trait, *θ*, and another parameter describing the strength of selection, *α*. In macroevolutionary applications, the parameter *α* is often translated into a characteristic time *t*_1/2_ = *ln*(2)/*α* needed for the trait to evolve halfway back to its optimum (Hansen, 1997). The heritability in POUMM is related to all three parameters of the Ornstein-Uhlenbeck process (Mitov and Stadler, 2016). We used the implementation by Mitov and Stadler (2017).

### 2.6 Testing for genetic associations with per-parasite pathogenicity

Previous studies identified a valine instead of an isoleucine at position 62 and 64 in the protease (I62V and I64V), and a proline instead of an alanine at position 272 in the reverse transcriptase as amino acid substitutions that could be associated with per-parasite pathogenicity (Ng *et al*., 2014) (see the Result section for more details). To test for associations of per-parasite pathogenicity with the substitutions I62V, I64V, and A272P, we regressed per-parasite pathogenicty, but also the other two traits, set-point viral load and CD4+ T cells decline, against categorical variables indicating the presence of the mutated amino acid at the respective position. The mutated amino acids were defined as a valine in position 62 and 64 of the protease, and a proline in position 272 of the reverse transcriptase. In these regressions, we also included cofactors such as age, sex and risk group.

An association with per-parasite pathogenicity was assessed in two ways. First, we regressed the proxy for per-parasite pathogenicity defined above — residuals from the age-adjusted tolerance curves — against the presence of the substitute amino acid. Second, we regressed the rate of change in the CD4+ T cells against the square of the logarithm to the base of 10 of the set-point viral load, including the presence of the mutated amino acid at the respective position as an interaction term. We also included the sex and age at infection as cofactors in the analysis:

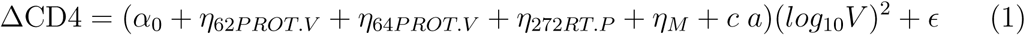

Here, ΔCD4 denotes the rate of change in the CD4+ T cell level per microliter blood per day. The parameter *α*_0_ describes the relationship between set-point viral load and CD4+ T cells decline for females with age zero that are infected with a virus that does not express any of the mutant amino acids we consider. The remaining parameters describe the offset that can be attributed to the various cofactors: *η*_62PROT.V_, *η*_64PROT.V_, and *η*_272RT.P_ describe the potential change in the relationship between set-point viral load and CD4+ T cell decline due to each considered amino acid substitution, and *η*_M_ quantifies the change due to being male. *c a* is the offset to *α*_0_ in an individual of age *a*. This procedure follows the approach we have adopted previously to test for the association of host factors with disease tolerance (Regoes *et al*., 2014).

## 3 Results

### 3.1 Heritability of set-point viral load confirmed

The heritability of the set-point viral load has previously been estimated from data of the Swiss HIV cohort study (Alizon *et al*., 2010) and from data of other cohorts (Hodcroft *et al*., 2014; Hollingsworth *et al*., 2010; Müller *et al*., 2011). The methods differed across these studies.

To test the conclusion of these studies, we applied donor-recipient regressions and the phylogenetic mixed models to the set-point viral loads from the 2014 individuals we included in the present study. The donor-recipient regressions were applied to 196 previously determined transmission pairs (Kouyos *et al*., 2014), while the phylogenetic methods were applied to a phylogenetic tree that was constructed from *pol* gene sequences (see Methods and Materials). Since set-point viral loads were significantly associated with sex, age at infection, and were higher in men who have sex with men, we also estimated the heritability of adjusted set-point viral loads as defined in the Methods and Materials.

Across all the methods we use, the estimates for heritability range from 8% to 29% (see Table 1, Fig 3B and Figure S2B. These estimates are all significantly larger than zero, except for the adjusted set-point viral load using the phylogenetic mixed-model that assumes neutral evolution. These results add to the growing consensus that set-point viral load is heritable.

**Table 1:** Heritability estimates for set-point viral load (spVL), CD4+ T cell decline (ΔCD4) and per-parasite pathogenicity (ppp) based on the phylogenetic mixed models assuming Brownian motion-type trait evolution (PMM) or trait evolution according to the Ornstein-Uhlenbeck process (POUMM). 95% confidence intervals are given in brackets.

|  | PMM | POUMM |
| --- | --- | --- |
| ΔCD4 (unadjusted) | 25% (9%–40%) | 17% (6%–29%) |
| ΔCD4 (adjusted) | 24% (7%–39%) | 17% (5%–30%) |
| spVL (unadjusted) | 12% (2%–28%) | 26% (8%–43%) |
| spVL (adjusted) | 8% (0%–26%) | 29% (12%–46%) |
| ppp | 22% (5%–39%) | 17% (4%–29%) |

**Figure 3:**
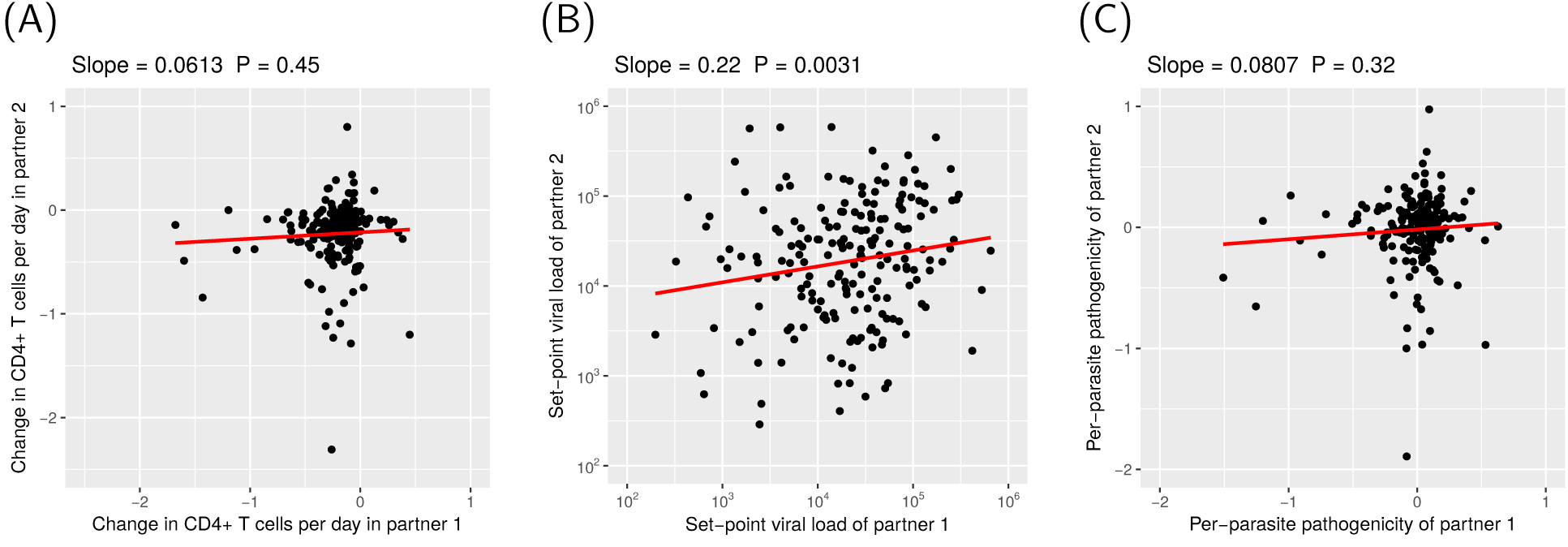
Heritability estimates from donor-recipient regressions. For these regressions we plotted the trait values for each partner (“partner 1” and “partner 2”) in the transmission pairs onto the x- and y-axis. Since we do not know the direction of the transmission in the pairs, the assignment of the partners to either x- or y-axis is random. (See also Figure S2.)

Interestingly, assuming stabilizing selection on the set-point viral load in our phylogenetic analysis led to a significantly better fit to the data than assuming neutral drift (Likelihood ratio test: *p* =1.2 × 10^−4^ for unadjusted and *p* = 8.8 × 10^−6^ for adjusted set-point viral loads). Thus, the estimates for the heritability of the set-point viral load with the best statistical support are 26% and 29% without and with adjustment for cofactors, respectively. Both of these estimates are significantly different from zero.

The optimal trait value *θ* of the Ornstein-Uhlenbeck process is estimated to be 10^4.0^ RNA copies per milliliter plasma for the unadjusted set-point viral load (95% CI: 10^1.6^ − 10^4.3^ RNA copies per milliliter plasma), very close to the mean of the set-point viral load in our study population 10^4.2^ RNA copies per milliliter plasma (see Table 2). The parameter measuring the strength of selection, *α* is high: 32.7 (95% CI: 0.03-57.6) and 39.4 (95% CI: 6.1-68.1) for unadjusted and adjusted set-point viral load, respectively (see Table 2). These parameter estimates are consistent with strong stabilizing selection around the current mean trait value.

**Table 2:** Estimates of the POUMM parameters related to selection for set-point viral load (spVL), CD4+ T cell decline (ΔCD4) and per-parasite pathogenicity (ppp) 95% confidence intervals are given in brackets.

| | $\alpha$ | $\theta$ | population mean |
| --- | --- | --- | --- |
| $\Delta\text{CD4}$ (unadjusted) | 4.1 (0.5 – 10.8) | -1.15 (-2.43 – -0.29) | -0.20 |
| $\Delta\text{CD4}$ (adjusted) | 3.8 (0.4 – 10.5) | -0.93 (-2.30 – -0.06) | -0.04 |
| spVL (unadjusted) | 32.7 (0.03 – 57.6) | 4.0 (1.6 – 4.3) | 4.2 |
| spVL (adjusted) | 39.4 (6.1 – 68.1) | -0.03 (-0.17 – 0.10) | -0.04 |
| ppp | 3.9 (0.5 – 10.4) | -0.89 (-2.17 – -0.09) | 0.00 |

### 3.2 Evidence for the heritability of CD4+ T cell decline

The set-point viral load is an important determinant of CD4+ T cell decline, and it is heritable. We therefore expect the CD4+ T cell decline to be also heritable “by association”. So far, however, there has been no evidence for the heritability of the CD4+ T cell decline or HIV virulence.

To assess the heritability of the CD4+ T cell decline, we applied the same methods as for the set-point viral load. Because the CD4+ T cell decline was associated significantly only with the age at infection we also conducted the analyses with age-adjusted CD4+ T cell declines as defined in the Methods and Materials.

The donor-recipient regression (Fig 3A) results in a hertability estimate, which is not significantly different from zero for both unadjusted and age-adjusted CD4+ T cell declines. This is likely the result of the low number of individuals in the transmission pairs (2 × 196), which limits the statistical power of the donor-recipient regressions.

Using the phylogenetic mixed models, however, we can incorporate all 2014 individuals of our study population, and obtain heritability estimates significantly larger than zero. Assuming neutral trait evolution (PMM) yields 25% and 24% for unadjusted and adjusted CD4+ T cell declines, respectively. With trait evolution according to the Ornstein-Uhlenbeck process we get 17% irrespective of any adjustment (see Table 1).

Again, assuming trait evolution according to an Ornstein-Uhlenbeck process has more statistical support than Brownian motion trait evolution (Likelihood ratio test: *p* = 6.9 × 10^−8^ for unadjusted and *p* = 2.6 × 10^−5^ for adjusted CD4+ T cell declines). But, unlike for the set-point viral load, the estimate of the optimal trait value *θ* (-1.15 per day, 95% CI: -2.43 to -0.29 per day) is significantly below the mean of the CD4+ T cell declines in our study population (-0.20 per day), and the estimated strength of selection *α* is an order of magnitude lower than that for the set-point viral load (4.1 and 3.8 for unadjusted and adjusted CD4+ T cell decline, respectively). See Table 2 for all selection related parameter estimates for this trait. These parameters characterize directional, rather than stabilizing selection, consistent with a slow time trend towards higher virulence.

### 3.3 Evidence for the heritability of the per-parasite pathogenicity

Lastly, we investigated if there is any evidence for the heritability of the per-parasite pathogenicity. The per-parasite pathogenicity is the component of virulence, which is determined by the pathogen genotype and independent of the pathogen load. Formally, we determine per-parasite pathogenicity by calculating the residual of the regression of the CD4+ T cell decline against the set-point viral load adjusted for the age of the infected individual (see Material and Methods and Fig 2).

A donor-recipient regression (Fig 3C) yields an estimate, which is not significantly different from zero, again likely due to the low number of transmission pairs. Using phylogenetic mixed models, we estimate a statistically significant heritability of 22% assuming Brownian motion-type trait evolution, and 17% for trait evolution according to the Ornstein-Uhlenbeck process (see Table 1).

As for the set-point viral load and the CD4+ T cell decline, assuming trait evolution governed by an Ornstein-Uhlenbeck process led to a significantly better fit than evolution according to Brownian motion (Likelihood ratio test: *p* = 2.2 × 10^−6^). The pattern of selection most consistent with the evolution of per-parasite pathogenicity is directional selection as for the CD4+ T cell decline: the optimal trait value is estimated to be significantly lower than the population mean, and the strength of selection is weak (see Table 2). This suggests that there is a slow time trend of increasing per-parasite pathogenicity.

### 3.4 Testing for genetic associations with per-parasite pathogenicity

It is known that HIV-1 subtype D has a higher per-parasite pathogenicity than subtype A. These intersubtype difference in disease progression have been mapped to the *pol* gene and were associated with replicative capacity (Ng *et al*., 2014). In particular, a valine instead of an isoleucine at position 62 and 64 in the protease (I62V and I64V), and a proline instead of an alanine at position 272 in the reverse transcriptase were found to be associated with high replicative capacity (Ng *et al*., 2014)

We tested if these amino acid polymorphisms are associated with set-point viral load, CD4+ T cell decline, and per-parasite pathogenicity within HIV-1 subtype B. Indeed isoleucine and valine at positions 62 and 64 of the protease, and alanine and proline at position 272 in the reverse transcriptase were the most prevalent amino acids in our study population. The set-point viral load was not associated with any of these polymorphisms. Before adjustment for multiple comparisons, the association of proline at position 272 in the reverse transcriptase with CD4+ T cell decline and per-parasite pathogenicity reach a significance level of *p* = 0.014 and *p* = 0.020, respectively, which is not significant after adjusting the p-values for three comparisons.

## 4 Discussion

In this study, we confirmed the heritability of the set-point viral load. Further, we report one of the first pieces of evidence for the heritability of the CD4+ T cell decline, a surrogate of virulence. We also found support for the hypothesis that the pathogen-load-independent virulence component is heritable. Lastly, the evolution of these three traits is significantly better described by the Ornstein-Uhlenbeck process than by Brownian motion.

Our study confirms previous studies that established the heritability of the set-point viral load in HIV infection (as reviewed by Müller *et al*. (2011) and Fraser *et al*. (2014)). In particular, our estimates are consistent with those from a donor-recipient regression in Hollingsworth *et al*. (2010), and two recent studies applying phylogenetic mixed models based on the Ornstein-Uhlenbeck process by Mitov and Stadler (2016) and Blanquart *et al*. (2017). Our analysis is also consistent with the study by Hodcroft *et al*. (2014) that reported a low heritability of 5.7% of the set-point viral load adjusted for covariates and assuming Brownian trait evolution. If we adjust for covariates and assume Brownian trait evolution, we obtain a heritability estimate of 8% that is not significantly different from zero. Assuming trait evolution according to the Ornstein-Uhlenbeck process, however, provides a significantly better fit to the adjusted set-point viral load data and yields a heritability estimate of 29%.

We find clear evidence for the heritability of the CD4+ T cell decline. As the level of CD4+ T cells is a defining characteristic of clinical AIDS, the CD4+ T cell decline is a good surrogate of virulence of HIV infection. Although the potential heritability of the rate of decline of CD4+ T cells has been investigated previously (Alizon *et al*., 2010), it was found to be not significantly different from zero. We attribute this discrepancy to the low sample size of the earlier study. In contrast to the 2014 individuals in our study population, Alizon *et al*. (2010) had enrolled only 1100 and investigated the heritability only in subpopulations consisting of a few hundered individuals.

The recent study by Blanquart *et al*. (2017) also reports heritability of the CD4+ T cell decline. For the HIV-1 subtype B, they estimate a heritability of 11% ranging from 0% to 19%. This estimate is within the 95% confidence intervals of our estimate, and our estimate of 17% is within the 95% confidence intervals of their estimate. Unlike our analysis, Blanquart *et al*. (2017) did not find support for the Ornstein-Uhlenbeck over the Brownian motion trait evolution model. Their estimate of the strength of selection on the CD4+ T cell decline is 0.095 whereas we estimate 3.8. This may be due to the lower sample size of 1170 individuals in the study by Blanquart *et al*. (2017). It may also be the result of differences in the inclusion criteria: Blanquart *et al*. (2017) included individuals with five CD4+ T cell measures between the time of their first positive HIV test and the beginning of treatment, while we require only three CD4+ T cells but in a more stringent time window that excludes the first 90 days after the estimated time of infection and time points, after the CD4+ T cell count fell below 100 cells per microliter blood. Differences in the tree reconstruction algorithm, however, are unlikely to explain the discrepancy. If we reconstruct the phylogenetic tree with RaxML (Stamatakis, 2006b), which is closely related to ExaML (Kozlov *et al*., 2015) that Blanquart *et al*. (2017) used, we obtain very similar parameter estimates (see Table S1). The heritability estimates based on the FastTree and RaxML trees are most discrepant for the adjusted set-point viral load when we use POUMM (29% versus 34% or 38%), but, because of the large confidence intervals, these estimates are not statistically different. With the RaxML trees, we also find that Ornstein-Uhlenbeck trait evolution has higher statistical support.

We also provide evidence for the heritability of the per-parasite pathogenicity. This trait describes the pathogenic potential of a viral strain that is independent of the load the strain attains in its host. We approximated this trait as the deviation of the CD4+ T cell decline observed in an individual from that predicted on the basis of the observed set-point viral load and the age of the infected host (see Methods). In addition to being determined by per-parasite pathogenicity, this deviation could be affected by further host factors, other sources of biological variation, and, of course, measurement noise, and should therefore be considered only as a surrogate for the per-parasite pathogenicity of a strain. It is important to note, however, that the uncertainties surrounding the quantification of the per-parasite pathogenicity make it more difficult to establish the heritability of this trait. The fact that we found evidence for heritability means that the signal in our surrogate measure of the per-parasite pathogenicity is not completely clouded by factors, for which we could not account.

For all three traits we considered, the Ornstein-Uhlenbeck trait evolution model has the best statistical support. For the set-point viral load the strength of selection is estimated to be high, and the inferred optimal value of the trait is close to the mean of the set-point viral load in our study population. Thus, set-point viral load is under significant stabilizing selection. The CD4+ T cell decline and the per-parasite pathogenicity, on the other hand, are not under strong stabilizing selection but directional selection — a scenario that the Ornstein-Uhlenbeck trait evolution model can capture with low selection strength *α* and an optimal trait value *θ* significantly different from the population mean. The parameter estimates of the Ornstein-Uhlenbeck model for these two traits are consistent with a slow but significant increase in HIV virulence over the past two decades (Pantazis *et al*., 2014). It is well-recognized that the Ornstein-Uhlenbeck trait evolution model can accommodate these two distinct patterns of selection. The difference in the nature of selection between set-point viral load and the other two traits is also the reason behind the bias in the heritability estimates based on Brownian motion trait evolution: the heritability of the set-point viral load is underestimated, while those of the other two traits are slightly overestimated. This conclusion is in agreement with simulation studies of this bias (Mitov and Stadler, 2017).

Intuitively, the heritability of the CD4+ T cell decline should be the combination of set-point viral load dependent and independent components of this trait. The heritability estimates we obtained conform surprisingly well with this expectation. The estimate of the heritability of the CD4+ T cell decline with the highest statistical support is 17%. The adjusted set-point viral load has a heritability of 29%. To approximate to what extent the heritability of the set-point viral load will trickle through to that of the CD4+ T cell decline, we need to factor in the correlation between these two traits. In our study population, the fraction of the variation in the CD4+ T cell decline explained by the set-point-viral load is *R*^2^ = 0.057, in very close agreement with estimates of this quantity in other study populations (Rodriguez *et al*., 2006). Thus, approximately 29% × 0.057 = 1.6% of the 17.4% of the heritability in the CD4+ T cell decline are due to the set-point viral load. The remainder — about 16% — should be independent of the set-point viral load. This agrees well with our estimate of the heritability of the per-parasite pathogenicity of 17%, especially given the uncertainty in all of these estimates.

The heritability of the per-parasite pathogenicity means that there are viral genes that influence the CD4+ T cell decline in ways that do not depend on the viral load. Generally, one conceivable such mechanism could be that viral genotypes with high per-parasite pathogenicity elicit ineffective immune responses that, rather than reducing viral load, accelerate CD4+ T cell decline. In terms of specific viral factors influencing per-parasite pathogenicity, studies that compare the disease course across HIV-1 subtypes are illuminating. In particular, the coreceptor usage (Daar *et al*., 2007) and *pol* replicative capacity (Barbour *et al*., 2004; Goetz *et al*., 2010; Ng *et al*., 2014) have been found to be associated with the rate of disease progression independently of the set-point viral load. Ng *et al*. (2014) identified three amino acid polymorphisms that are associated with replicative capacity. We tested if these polymorphisms are associated with per-parasite pathogenicity, CD4+ T cell decline and set-point viral load. While we could not establish any clear-cut association after adjusting for multiple comparisons, a proline at position 272 of the reverse transcriptase was the most promising candidate. The sample size for this test was lower (n=1222) than the size of our study population (n=2014) because the sequence region containing position 272 in the reverse transcriptase was not available in all individuals. This polymorphisms should be tested in the future for an association with CD4+ T cell decline and per-parasite pathogenicity in a larger study population.

Previously, Pagel’s λ was also employed to estimate the heritability of the set-point viral load from a phylogenetic tree of HIV *pol* sequences (Alizon *et al*., 2010). Leventhal and Bonhoeffer (Leventhal and Bonhoeffer, 2016), however, have argued recently that Pagel’s λ implicitly assumes the trees to be ultrametric. Thus, for non-ultrametric phylogenetic trees, such as the one we analyzed, this essential assumption of Pagel’s λ is violated, and this method should therefore not be used. We nevertheless provide heritability estimates for comparison in Figure S1. With Pagel’s λ the set-point viral load and the CD4+ T cell decline are also found to be significantly heritable.

The heritability of traits from donor to recipient is sometimes interpreted as the quantitative measure of the extent to which the trait is “under the control of the virus”. It has been pointed out that the viral genome carries an imprint of past environments, in particular the immune responses experienced in former hosts (Bartha *et al*., 2013, 2017; Carlson *et al*., 2014, 2016; van Dorp *et al*., 2014). Thus, separating virus from host effects is challenging. What is undisputed is that the trait in the new host is in part encoded in the viral genome. Information on associations with viral genes is very valuable, especially for traits that cannot be determined instantaneously, such as the CD4+ T cell decline.

In summary, we presented a comprehensive evolutionary analysis of the components of HIV virulence. We established that viral load dependent and independent virulence components, as well as overall virulence are heritable. This strongly suggests that these virulence components are, at least in part, encoded in the viral genome. Future research will need to identify the specific genetic polymorphisms associated with these virulence components.

## 5 Acknowledgments

We are grateful to Oliver Laeyendecker for making us aware of the amino acid polymorphisms that could be associated with per-parasite pathogenicity.

We thank the patients who participate in the Swiss HIV Cohort Study (SHCS); the physicians and study nurses, for excellent patient care; the resistance laboratories, for high-quality genotypic drug resistance testing; SmartGene (Zug, Switzerland), for technical support; Brigitte Remy, RN, Martin Rickenbach, MD, Franziska Schöni-Affolter, MD, and Yannick Vallet, MSc, from the SHCS Data Center (Lausanne, Switzerland), for data management; and Danièle Perraudin and Mirjam Minichiello for administrative assistance.

RRR acknowledges the financial support of the Swiss National Science Foundation (grant number: 31003A_149769). RDK and AM have been supported by the Swiss National Science Foundation (Grant number: BSSGI0_155851). This study has been performed within the framework of the Swiss HIV Cohort Study, supported by the Swiss National Science Foundation (grant number 33CS30_148522) and was further supported by the SHCS research foundation. The Swiss HIV Drug Resistance database is supported by SNF project 320030_159868 to H. F. G, the Yvonne-Jacob Foundation; Gilead, Switzerland (1 unrestricted grant to the SHCS Research Foundation and 1 unrestricted grant to H. F. G.); and the University of Zurich’s Clinical Research Priority Program (Viral Infectious Diseases: Zurich Primary HIV Infection Study; to H. F. G.)

The data are gathered by the 5 Swiss University Hospitals, 2 Cantonal Hospitals, 15 affiliated hospitals and 36 private physicians (listed in http://www.shcs.ch/31-health-care-providers). The members of the Swiss HIV Cohort Study are: Aubert V, Barth J, Battegay M, Bernasconi E, Böni J, Bucher HC, Burton-Jeangros C, Calmy A, Cavassini M, Egger M, Elzi L, Fehr J, Fellay J, Furrer H (Chairman of the Clinical and Laboratory Committee), Fux CA, Gorgievski M, Günthard H (President of the SHCS), Haerry D (deputy of “Positive Council”), Hasse B, Hirsch HH, Hösli I, Kahlert C, Kaiser L, Keiser O, Klimkait T, Kouyos R, Kovari H, Ledergerber B, Martinetti G, Martinez de Tejada B, Metzner K, Müller N, Nadal D, Pantaleo G, Rauch A (Chairman of the Scientific Board), Regenass S, Rickenbach M (Head of Data Center), Rudin C (Chairman of the Mother & Child Substudy), Schöni-Affolter F, Schmid P, Schultze D, Schüpbach J, Speck R, Staehelin C, Tarr P, Telenti A, Trkola A, Vernazza P, Weber R, Yerly S.

## Supplementary Figures and Tables

**Figure S1:**
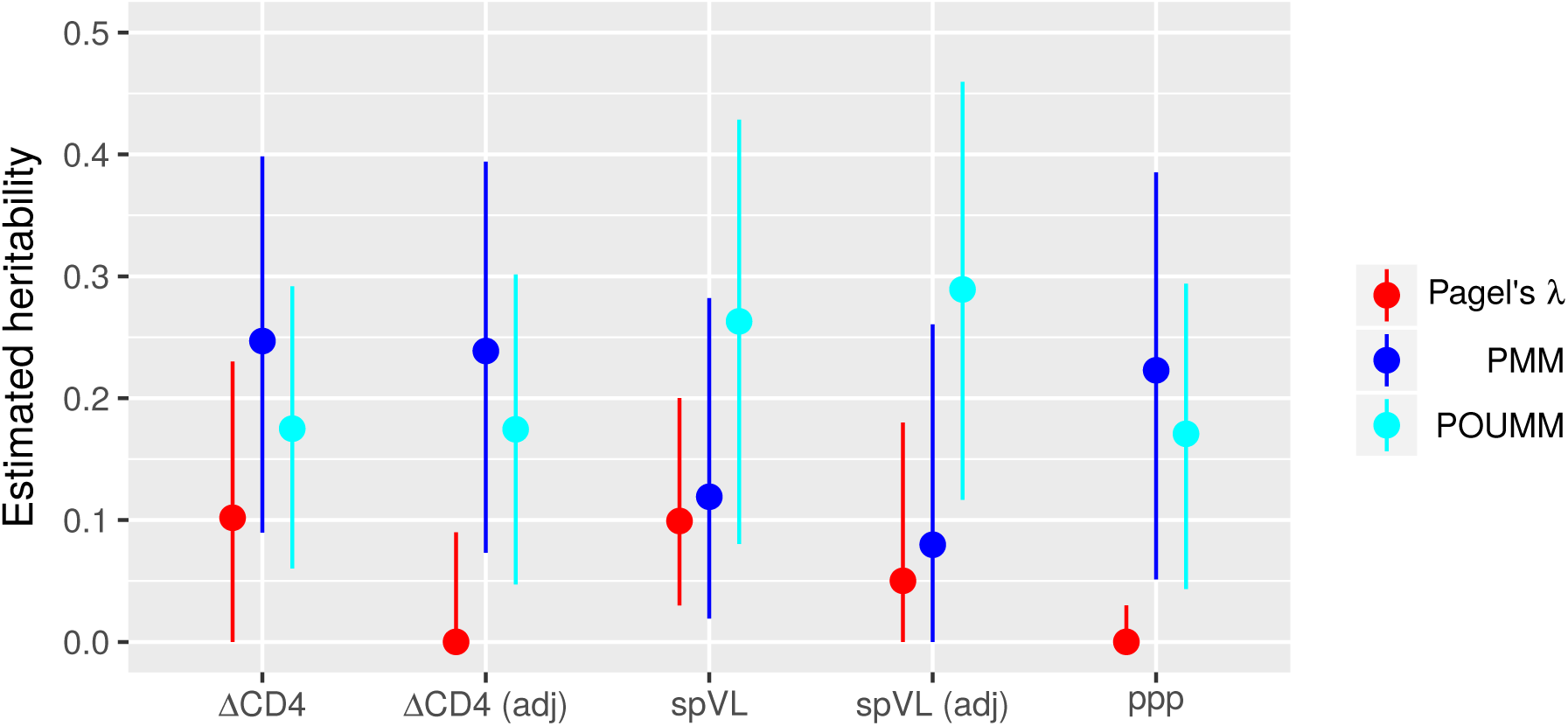
Heritability estimates using using Pagel’s λ in comparison to those using PMM and POUMM. The points are the point estimates, the vertical lines show the 95% confidence intervals. We show these estimates only for comparison with older papers that used Pagel’s λ on unadjusted traits. Because of the non-ultrametric nature of our phylogenetic tree Pagel’s λ is not appropriate to use.

**Figure S2:**
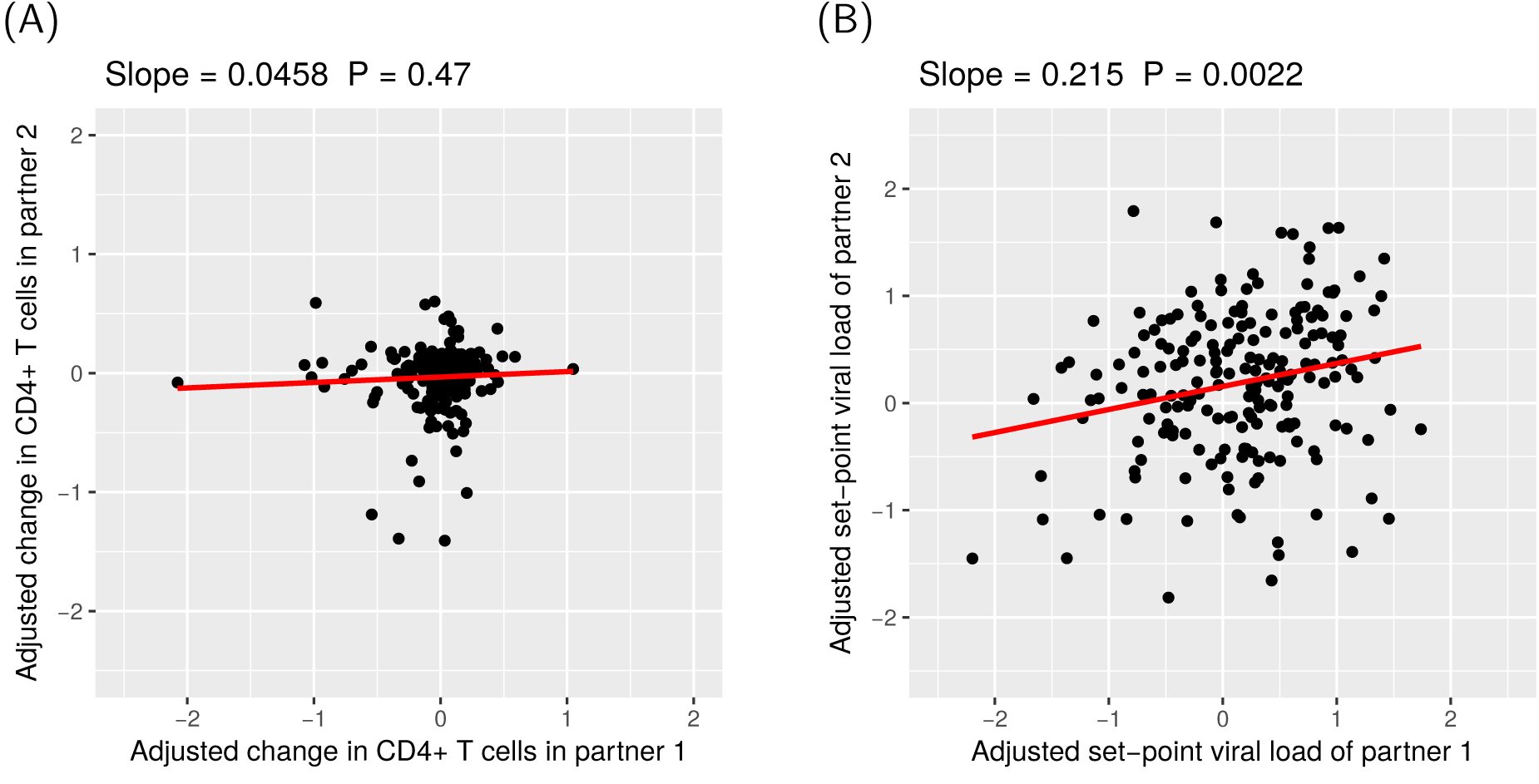
Heritability estimates from donor-recipient regressions on adjusted CD4+ T cell decline and set-point viral loads. For the definition of the adjusted trait values see the Methods and Materials. (See also Fig 3.)

**Table S1:** Comparison of the heritability estimates using different tree reconstruction algorithms. For all traits, POUMM fits the data significantly better than PMM.

| Method | Tree reconstruction | ΔCD4 (unadjusted) | ΔCD4 (adjusted) | spVL (unadjusted) | spVL (adjusted) | ppp |
| --- | --- | --- | --- | --- | --- | --- |
| PMM (ML) | FastTree | 25% (9%–40%) | 24% (7%–39%) | 12% (2%–28%) | 8% (0%–26%) | 22% (5%–39%) |
|  | RaxML-CAT | 24% (9%–39%) | 0% (0%–10%) | 13% (3%–29%) | 0% (0%–12%) | 23% (7%–39%) |
|  | RaxML-Gamma | 24% (9%–39%) | 0% (0%–3%) | 13% (3%–29%) | 11% (2%–25%) | 23% (6%–39%) |
| PMM (MCMC) | FastTree | 24% (12%–36%) | 23% (9%–36%) | 13% (4%–23%) | 9% (0%–19%) | 21% (9%–32%) |
|  | RaxML-CAT | 24% (13%–34%) | 25% (14%–34%) | 14% (5%–24%) | 9% (0%–20%) | 24% (13%–35%) |
|  | RaxML-Gamma | 24% (13%–35%) | 24% (14%–36%) | 14% (5%–25%) | 11% (1%–22%) | 22% (6%–33%) |
| POUMM | FastTree | 17% (6%–29%) | 17% (5%–30%) | 26% (8%–43%) | 29% (12%–46%) | 17% (4%–29%) |
|  | RaxML-CAT | 20% (8%–30%) | 21% (9%–32%) | 33% (19%–48%) | 38% (22%–52%) | 19% (7%–31%) |
|  | RaxML-Gamma | 19% (8%–30%) | 20% (8%–33%) | 30% (15%–46%) | 34% (18%–50%) | 19% (6%–31%) |

